# Comparative Systematic Analysis of Gray Matter Biophysical Models on a public dataset

**DOI:** 10.1101/2025.06.19.660512

**Authors:** Santiago Mezzano, Quentin Uhl, Tommaso Pavan, Jasmine Nguyen-Duc, Hansol Lee, Susie Huang, Ileana Jelescu

## Abstract

Biophysical models of diffusion tailored to characterize gray matter (GM) microstructure are gaining traction in the neuroimaging community. NEXI, SMEX, SANDI, and SANDIX represent recent efforts to incorporate different microstructural features,such as soma contributions and inter-compartment exchange, into the diffusion MRI (dMRI) signal. In this work, we present a comparative evaluation of these four gray matter models on a single, publicly available in vivo human dataset, the Connectome Diffusion Microstructure Dataset (CDMD), acquired with two diffusion times. Using the open-source Gray Matter Swiss Knife toolbox, we estimate cortical microstructure metrics in 26 healthy subjects and evaluate goodness of fit, anatomical patterns and consistency with previous studies.

CDMD data yielded GM parameter estimates consistent with values reported in previous studies. This retrospective cross-model analysis establishes the feasibility of estimating exchange models from only two diffusion times and highlights trade-offs in biological specificity, model complexity, and fitting robustness, critical considerations when choosing a model for future clinical and research applications.

## 1 Introduction

Diffusion-weighted magnetic resonance imaging (dMRI) is at the forefront of neuroimaging technologies, offering a unique window into the microstructural intricacies of brain tissue and cellular structures in vivo, non-invasively [1]. dMRI exploits the micrometer-scale movement of water molecules, whose diffusion patterns are influenced by cellular geometry, membrane permeability, and the presence of intracellular organelles at comparable scales [2, 3]. This holds promise for both advancing our understanding of brain structure and enabling the early detection of neurological and psychiatric disorders [4, 5].

The sensitivity of the dMRI signal to tissue microstructure is indirect, and cellular parameters can be quantified through biophysical modeling, which links biomedical imaging with fundamental physics paradigms [6]. Biophysical models are, however, tissue specific.

In recent years, the dMRI community has taken up the challenge to develop and validate novel biophysical models tailored to brain gray matter (GM), to account for its specificities in the form of a permeable neurite membrane [7, 8] (unlike myelinated impermeable axons in white matter) or a non-negligible density of cell bodies [9].

By quantifying the microstructural features of the human cortex in vivo, and interpreting their neurobiological signature, GM biophysical models hold the promise to significantly advance our capabilities in both research and clinical settings.

This comparative analysis will thus focus on four biophysical models of GM, all implemented as part of an open-source tool named Gray Matter Swiss Knife [10]. The GM models under consideration are NEXI [7], SMEX [8], SANDI [9], and SANDIX [8]. As these models make their way towards translation to clinical MRI scanners [11, 12], it is important to test their performance on publicly available datasets, more of which will hopefully be acquired and shared in the years to come. Here, we performed a retrospective analysis on 26 healthy subjects from the MGH Connectome Diffusion Microstructure open-source Dataset (CDMD) [13], leveraging its ideal acquisition parameters of multi-diffusion times and high gradient strengths required for estimating these gray matter models [7, 9, 8].

The goal of this study is to: (i) evaluate the feasibility and reliability of estimating each of the four models on a diffusion protocol comprising two diffusion times and multiple shells, (ii) quantify and compare cortical Region of Interest (ROI) values and surface maps with previous literature on these models based on dedicated and more comprehensive datasets. Moreover, NEXI has only been estimated so far in the human brain using dMRI protocols with three [14] or four [15, 12] diffusion times. The feasibility of NEXI/SMEX estimation based on two diffusion times, previously shown in ex vivo mouse brain [8], would lower the minimum acquisition time and thus increase its potential for clinical translation.

## 2 Theory

Here we briefly describe the assumptions behind each of the GM models under consideration (Figure 1). More details can be found in the original papers introducing them [7, 9, 8].

**Figure 1.**
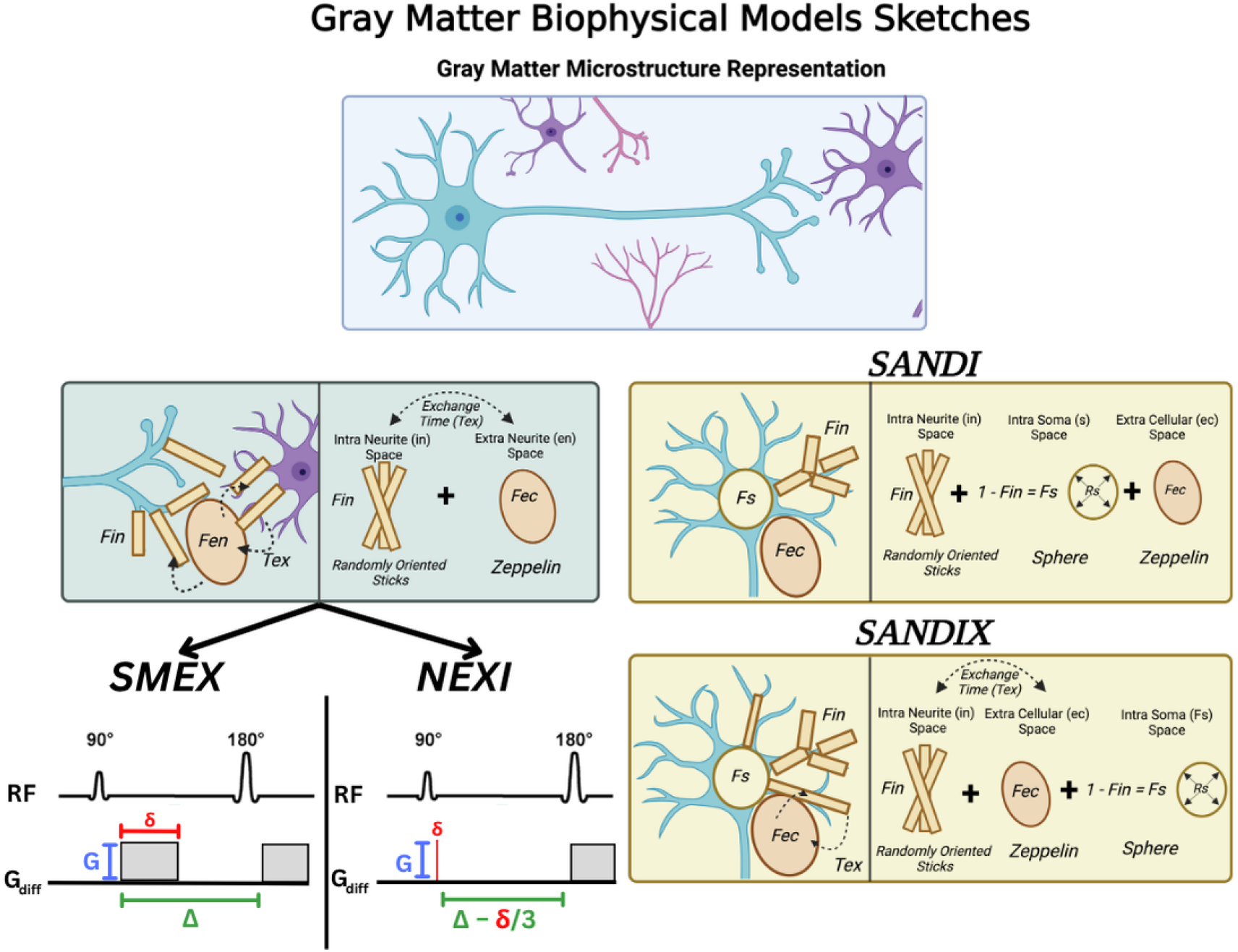
Graphical summary of four gray matter biophysical models—NEXI, SMEX, SANDI, and SANDIX—used to model the diffusion MRI (dMRI) signal in the cortex. Each model includes different compartments to represent intra-neurite space, intra-soma space, extra-neurite space and water exchange mechanisms. For NEXI and SMEX we illustrate the corresponding PGSE (Pulsed Gradient Spin Echo) sequence which highlights their assumptions about diffusion time (*t*_*d*_), with SMEX accounting for the gradient duration *δ* and NEXI incorporating the correction *t*_*d*_ = Δ − *δ/*3 to account for finite gradient durations.

### 2.1 Incorporating the Soma compartment - *SANDI*

Soma and Neurite Density Imaging (SANDI) [9] is a biophysical model that introduces a three-compartment framework representing:

- **Neurites:** Randomly oriented sticks with uni-dimensional intra-stick axial diffusivity (*D*_*i*,||_) and a relative signal fraction (*f*).
- **Soma:** Spheres with an apparent radius (*R*_*s*_), fixed intra-sphere diffusivity (*D*_*s*_), and a relative signal fraction (*F*_*s*_). Diffusion in spheres relies on the Gaussian Phase Approximation.
- **Extracellular Space:** Modeled with Gaussian isotropic diffusion (*D*_*e*_) and a relative signal fraction (1 − *f* − *F*_*s*_).

SANDI thus allows for the independent estimation of five parameters: neurite signal fraction (*f*), soma signal fraction (*F*_*s*_), intra-stick diffusivity (*D*_*i*,||_), extracellular diffusivity (*D*_*e*_), and soma radius (*R*_*s*_) (Figure 1).

This model does not account for inter-compartment exchange, hence the need to use a protocol with short diffusion time (Δ ≤ 20 ms) to keep the assumption of impermeable stick compartments valid. High b-values (e.g. above *b* = 6 ms*/µ*m^2^) within short diffusion times can only be acquired by systems with ultra-strong gradients, such as preclinical scanners or new generation dedicated human scanners (Connectom 1.0, Connectom 2.0, Magnus). Longer diffusion times would cause the exchange between compartments to contaminate the signal and affect the accuracy of the estimated parameters.

Recent studies have tested the effectiveness of SANDI in multiple contexts. Initially, the model was shown to produce soma and neurite signal fraction maps using multi-shell sequences with *b*-values up to 10 ms/*µ*m^2^ on a human Connectom scanner, which agreed with histological images of brain architecture [9]. Additionally, parameter stability was confirmed in *in vivo* healthy mouse brains, with strong correlations between SANDI-derived soma signal fractions and histology of cell density data from the Allen Brain Atlas [16]. Recent research reported SANDI estimates across the GM cortical ribbon in the human brain using a 3T scanner with weaker gradients and a lower maximum *b*-value of 6 ms/mm^2^ [11], albeit at a longer diffusion time (Δ = 39.07 ms), which likely violates the impermeability assumption.

**Created in BioRender**. Mezzano, S. (2025) https://BioRender.com/q54w840 **Abbreviations:** *f*_neurite_ (*F*_in_): intra-neurite signal fraction; *f*_extracellular_ (*F*_ec_): extra-neurite signal fraction; *f*_soma_ (*F*_s_): Intra Soma Space; *R*_soma_ (*R*_s_): apparent soma radius; *t*_ex_: exchange time.

### 2.2 Accounting for Exchange - *NEXI, SMEX*

Simulations and experimental data have consistently shown that water exchange between neurites and the extracellular space in GM is non-negligible over typical diffusion times of MRI experiments (20–100 ms) as most neurites are unmyelinated and thus permeable [17, 7, 8, 18].

To address this challenge, two signal models were proposed in parallel, Neurite Exchange Imaging (NEXI) [7, 15] and the Standard Model with EXchange (SMEX) [8, 12], which are based on the same underlying microstructure but differ in their treatment of the diffusion sequence: NEXI assumes the narrow pulse approximation (NPA), whereas SMEX does not. By incorporating an exchange parameter, NEXI and SMEX eliminate the requirement for short diffusion times and can thus be implemented on clinical-grade scanners. Both models are parametrized by four parameters to be estimated (Figure 1).

- **Neurites:** Same as SANDI; modeled as a collection of randomly oriented sticks, with intra-stick uniaxial diffusivity (*D*_*i*,||_) and a relative signal fraction (*f*).
- **Extracellular Space:** Same as SANDI; considered Gaussian isotropic with characteristic diffusivity (*D*_*e*_).
- **Exchange Process:** Exchange between neurites and extracellular space with a characteristic exchange time (*t*_*ex*_).

The signal from the soma is not explicitly considered in NEXI or SMEX and is assumed to be pooled with the extra-cellular space [7].

While NEXI assumes short gradient pulse durations *δ* under the NPA, SMEX additionally accounts for finite, and potentially long, pulse durations, making it ideally suited for realistic acquisition protocols on clinical systems. Indeed, NEXI, based on the Kärger model approximation [19], simplifies exchange dynamics by modeling the Pulsed Gradient Spin Echo (PGSE) diffusion encoding pulses as instantaneous Dirac pulses separated by a duration Δ. This assumption allows a more straightforward solution, at the cost of some accuracy. Note however that defining the diffusion time as *t*_*d*_ = Δ − *δ/*3 allows for some correction for the finite pulse duration *δ*. SMEX on the other hand uses numerical integration and accounts for dynamics during the diffusion gradient pulse as well.

One requirement shared by both NEXI and SMEX is that of a protocol that samples multiple diffusion times *t*_*d*_ or Δ in order to robustly estimate exchange. This requirement substantially increases the acquisition time.

NEXI has been tested on both preclinical and human scanners to quantify GM microstructure [7, 15]. The first NEXI study in the human cortex found very good scan-rescan reproducibility while maintaining subject-specific GM variability [15]. This work also emphasized the importance of accounting for the Rician noise floor when fitting magnitude data, which was added to the model termed NEXI_RM_ (for Rician Mean).

SMEX was initially implemented and tested on preclinical data in ex vivo mouse cortex[8]. Recently, the performance of SMEX vs NEXI was also demonstrated on a clinical scanner with longer gradient pulses [12].

### 2.3 Accounting for Soma and Exchange - *SANDIX*

A combination of SANDI and NEXI is Soma And Neurite Density Imaging with Exchange (SANDIX) [8]. SANDIX accounts for an impermeable soma compartment in addition to the exchanging neurites and extra-cellular space. This results in a three-compartment model with six free parameters: *f*, *F*_*s*_, *R*_*s*_, *D*_*i*_, *D*_*e*_, and *t*_ex_ (Figure 1). The feasibility of estimating so many parameters has only been tested on an extensive ex vivo preclinical dataset [8], and has yet to be evaluated in the human cortex.

As mentioned, all previous human in vivo studies estimated NEXI from data parsing four diffusion times. SMEX and SANDIX were previously estimated in the mouse cortex using two diffusion times on a pre-clinical scanner [8]. The MGH Connectome Diffusion Microstructure Dataset (CDMD) [13] used in the current study covers a larger cohort size than previous NEXI/SMEX datasets, and samples only two diffusion times, making it the first instance where these models are applied with this type of reduced acquisition in the human cortex. In this context, the value of this dataset lies in the fact that the acquisition protocol was not optimized for any specific model, allowing for an unbiased comparison across modeling frameworks.

## 3 Methods

We performed a retrospective analysis on 26 healthy young subjects downloaded from the CDMD [13], to compare cortical microstructure estimates stemming from each of the GM models. We also conducted a post-hoc analysis comparing NEXI and SMEX to elucidate further differences in quality of fit estimates and model behavior in this dataset.

### 3.1 Participants

The study protocol was approved by the institutional review board of Mass General Brigham, following compliance with all relevant ethical regulations. Written informed consent was obtained from all participants. Data were acquired from 26 healthy, cognitively normal adults (36.8±14.6 years old, 17 females).

### 3.2 Connectome Diffusion Microstructure Dataset (CDMD)

Detailed methods can be found in [13]. Briefly, data were acquired on a 3 T Connectom MRI scanner, with a gradient coil of 300 mT/m. For anatomical reference, T1 images were acquired using a multiecho magnetization-prepared gradient echo (MEMPRAGE) sequence, at 1-mm isotropic resolution. Diffusion-weighted images were acquired using a single-refocused pulsed-gradient spin-echo echo-planar imaging (EPI) sequence, with *b*-values ranging from 0.05 to 6 ms/mm^2^ at Δ = 19 *ms* and from 0.2 to 17.8 ms/mm^2^ at Δ = 49 *ms*, in 16 increments. This included 32 or 64 diffusion encoding directions depending on the *b*-value, alongside 50 non-diffusion-weighted (*b* = 0) images. Other fixed parameters were gradient duration *δ* of 8 ms, TE/TR of 77 ms/3.8 s, at 2-mm isotropic resolution. The total scan time for the diffusion MRI protocol was 55 minutes. Downloaded diffusion data were already pre-processed with a standard pipeline, correcting for gradient nonlinearity, susceptibility and eddy current-induced image distortions, and motion [13]. The full acquisition protocol can be found in the Supplementary Material (Figure **??**).

Importantly, for a subset of 14 participants, phase images in addition to magnitude images were available. From complex-valued data, real-valued diffusion data were extracted after filtering the phase, as described by [13]. This method preserved the Gaussian noise distribution in the data which better aligns with the assumptions underlying MP-PCA denoising. Additionally, the elevated Rician noise floor at high b-values in the magnitude data was largely attenuated in the real-valued data. Removing or accounting for the Rician noise floor has been shown to be critical when processing gray matter biophysical models, particularly when estimating parameters related to exchange between compartments [15].

Here, we processed both magnitude data from 26 subjects as well as real-valued data from 14 subjects to investigate accuracy in parameter estimates within and between biophysical models across varying de-noising strategies. However, a more thorough analysis and comparison of models was done on the 26 subjects with magnitude data, leveraging the increased number of subjects and the more typical context of magnitude images in clinical settings.

### 3.3 Gray Matter Swiss Knife

Gray Matter Swiss Knife is an open source python package [10] that implements the estimation of various parameter maps from four different gray matter biophysical models (SMEX, NEXI, SANDI, SANDIX). Gray Matter Swiss Knife uses nonlinear least squares (NLS) with initial grid search, utilizing the L-BFGS-B algorithm and minimize function from the scipy library in Python [20].

For datasets where both real-valued and magnitude data were available (N=14), NEXI and SMEX were estimated without Rician mean correction in real-valued data, and with Rician mean correction (RM) in the magnitude data. Estimates were compared.

For all magnitude datasets (N=26), the four models were estimated. The noise map calculated from MP-PCA denoising at high SNR (low b-values) was used to correct for Rician Mean [15] in NEXI, SMEX, SANDIX and SANDI.

Finally, as SANDI assumes no exchange between compartments and is not typically applied on multiple diffusion time data, model estimates were additionally calculated for the shorter diffusion time Δ = 19 ms only. The shorter diffusion time was chosen to evaluate the performance of SANDI in the best possible conditions for this model, when the signature of exchange is limited.

### 3.4 Cortical Parcellation, Volumetric Maps and Surface Maps

GM ROIs from the Desikan-Killiany-Tourville (DKT) atlas [21] were segmented on the anatomical MEMPRAGE image using FastSurfer [22] and transformed into native diffusion space using linear registration of distortion-corrected b = 0 images to T1 images. The cortical ribbon was segmented by merging the gray matter ROIs obtained from the DKT segmentation. Mean parameter estimates were calculated by finding the median in every DKT ROI and then calculating the mean from all ROIs in the cortical ribbon. Calculating the median as opposed to the mean at the ROI level reduces the impact of outliers, which can arise due to noise, artefacts, partial volume or segmentation errors.

It also reproduces the same analysis that other papers have used to quantify average gray matter microstructure [15, 11].

To visually inspect the parameter maps generated for each biophysical model, average maps from the 26 subjects were generated in standard space (MNI) [23].

Moreover, SMEX, NEXI parameter maps were transformed into Freesurfer’s fsaverage subject space and projected onto cortical surfaces to visualize and compare anatomical trends with those presented in previous works [15, 12, 24]. One subject was discarded from surface map reconstruction due to an incomplete cortical coverage within the field of view in its CDMD dataset.

### 3.5 Quality of Fit

In order to compare the quality of the fit of the model on the measured data, we compared the Measured vs Fitted Signal as a function of 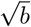 at different diffusion times, and also computed the corrected Akaike Information Criterion (AICc) [25] for each model. This metric evaluates the goodness of fit while penalizing model complexity, aiming to achieve a balance between accuracy and simplicity.

### 3.6 Simulations

Additionally, we report SMEX and NEXI estimates against ground truth values. Synthetic SMEX signals were generated using the same values of *δ*, diffusion times, and b-values as in the experimental acquisition.

The ground truth parameters for each signal were randomly sampled from the combined NEXI and SMEX experimental parameter estimates, under the constraint that *D*_*i*_ *> D*_*e*_ [26, 27, 28].

For each powder-averaged signal at a given *b*-*t*_*d*_ combination, we generated a number of Rician noise realizations equal to the number of diffusion directions acquired for that specific *b*-*δ*, for each ground truth instance. The signal-to-noise ratio (SNR) for each ground truth instance was randomly drawn from the in vivo SNR distribution at *b* = 0 and *b* = 1 ms*/µ*m^2^.

The noisy signals were then averaged to emulate powder-averaged magnitude images, which improves the SNR but does not reduce the Rician noise floor. In total, a dataset of 10’000 ground truth parameter combinations was constructed using this approach.

The generated signals were then fitted using both the SMEX and NEXI models to assess error propagation when the ground truth perfectly follows the SMEX model. The discrepancy between the ground truth and the estimated parameters was quantified. A density-colored scatter plot was used to visualize the spread of these estimation errors.

## 4 Results

### 4.1 Quantitative Comparison Across All Gray Matter Biophysical Models

#### 4.1.1 Summary of Parameter Estimates

We first present the parameter estimates of the cortical ribbon derived from the real-valued data of 14 subjects (Table 1, section A) and the magnitude data of 26 subjects (Table 1, section B), along with their 95% confidence intervals. In addition, we provide mean parameter estimates from six previous studies examining cortical ribbon metrics derived from SANDI [11, 29] and NEXI/SMEX [15, 12, 14] for comparison (Table 1, section C).

**Table 1:** Mean estimates and 95% confidence intervals for biophysical models.

| A: Real-valued data from 14 subjects |  |  |  |  |  |  |
| --- | --- | --- | --- | --- | --- | --- |
| MODEL | $t_{\text{ex}}$ (ms) | $D_i$ ($\mu\text{m}^2/\text{ms}$) | $D_e$ ($\mu\text{m}^2/\text{ms}$) | $f$ | $F_s$ | $R_s$ ($\mu\text{m}$) |
| NEXI | 68.8 (25.9, 150.0) | 2.94 (0.89, 3.50) | 0.90 (0.80, 1.12) | 0.30 (0.13, 0.42) | - | - |
| SMEX | 41.4 (18.5, 125.2) | 2.38 (1.33, 2.99) | 0.99 (0.87, 1.23) | 0.36 (0.28, 0.45) | - | - |
| SANDI | - | 2.90 (0.80, 3.50) | 0.79 (0.69, 0.87) | 0.19 (0.07, 0.26) | 0.09 (0.01, 0.24) | 24.3 (12.1, 30.0) |
| SANDIX | 40.1 (25.9, 105.4) | 2.37 (1.83, 2.68) | 1.03 (0.74, 1.38) | 0.32 (0.28, 0.37) | 0.10 (0.05, 0.21) | 17.5 (12.7, 22.1) |
| B: Magnitude Data from 26 subjects |  |  |  |  |  |  |
| MODEL | $t_{\text{ex}}$ (ms) | $D_i$ ($\mu\text{m}^2/\text{ms}$) | $D_e$ ($\mu\text{m}^2/\text{ms}$) | $f$ | $F_s$ | $R_s$ ($\mu\text{m}$) |
| NEXI<sub>RM</sub> | 29.3 (3.96, 125.4) | 2.97 (0.78, 3.50) | 0.95 (0.80, 1.34) | 0.38 (0.17, 0.55) | - | - |
| SMEX<sub>RM</sub> | 21.1 (12.7, 49.9) | 2.46 (1.28, 3.03) | 1.00 (0.85, 1.31) | 0.37 (0.25, 0.48) | - | - |
| SANDI | - | 3.10 (1.23, 3.50) | 0.78 (0.68, 0.87) | 0.20 (0.08, 0.27) | 0.10 (0.01, 0.25) | 24.3 (13.06, 30.00) |
| SANDI ($\Delta = 19$ ms only) | - | 2.92 (1.37, 4.47) | 0.90 (0.23, 1.57) | 0.18 (0.09, 0.27) | 0.16 (0.06, 0.26) | 9.5 (7.31, 11.69) |
| SANDIX | 25.9 (20.5, 32.9) | 2.20 (1.60, 2.61) | 1.09 (0.76, 1.60) | 0.30 (0.22, 0.36) | 0.09 (0.03, 0.21) | 17.2 (12.9, 22.1) |
| C. Previous Papers |  |  |  |  |  |  |
| MODEL | $t_{\text{ex}}$ (ms) | $D_i$ ($\mu\text{m}^2/\text{ms}$) | $D_e$ ($\mu\text{m}^2/\text{ms}$) | $f$ | $F_s$ | $R_s$ ($\mu\text{m}$) |
| NEXI<sub>RM</sub> CONNECTOME 1 (Uhl, 2024) | 42.3 [40.0, 44.7] | 3.35 [3.32, 3.38] | 0.92 [0.91, 0.93] | 0.38 [0.37, 0.38] | - | - |
| NEXI<sub>RM</sub> (Uhl, 2025) | 41.1 [15.5, 66.7] | 2.61 [2.03, 3.20] | 1.59 [1.18, 2.00] | 0.51 [0.41, 0.61] | - | - |
| SMEX<sub>RM</sub> (Uhl, 2025) | 36.8 [17.6, 58.0] | 2.45 [1.99, 2.90] | 1.23 [0.88, 1.61] | 0.42 [0.34, 0.50] | - | - |
| NEXI CONNECTOME 1 (Chan, 2025) | 44.4 [30.2, 77.6] | 3.00 [3.00, 3.00] | 0.89 [0.81, 0.95] | 0.28 [0.27, 0.30] | - | - |
| NEXI CONNECTOME 2 (Chan, 2025) | 20.8 [14.9, 33.5] | 3.00 [3.00, 3.00] | 0.88 [0.81, 0.94] | 0.34 [0.32, 0.34] | - | - |
| SANDI ($\Delta = 39.07$ ms) (Schiavi, 2023) | - | 2.45 [2.35, 2.55] | 1.36 [1.27, 1.45] | 0.34 [0.32, 0.36] | 0.26 [0.24, 0.29] | 12.7 [12.3, 13.1] |
| SANDI ($\Delta = 19$ ms) (Lee, 2024) | - | 1.83 [1.53, 2.13] | 1.60 [1.53, 1.67] | 0.42 [0.40, 0.44] | 0.17 [0.16, 0.18] | 9.26 [9.11, 9.41] |

Table 1 shows that parameter estimates across models are fairly consistent between denoising strategies (complex vs magnitude denoising) and with previous literature reports using other datasets. Nevertheless, within the CDMD dataset, the parameter that varies the most between real- vs magnitude data is the exchange time (*t*_*ex*_), with consistently higher values in real-valued vs magnitude data (*t*_ex_ = 68.8 ms vs 29.3 ms for NEXI, and *t*_ex_ = 41.4 ms vs 21.1 ms for SMEX). SANDIX also yielded similar differences between real-valued (*t*_ex_ = 40.1 ms) and magnitude (*t*_ex_ = 25.9 ms) data. All NEXI estimates on real-valued data yielded similar cortical *t*_*ex*_ between 40 and 70 ms, which is aligned with previous Connectom 1 estimates [15] but less with recent Connectom 2 results [14]. NEXI estimates on magnitude data with Rician Mean correction yielded lower exchange time estimates (20 — 30 ms) than previous similar implementations (36 — 42 ms).

For intra-cellular diffusivity, all models show similar values for real and magnitude data. The NEXI and SANDI estimates are higher than SMEX and SANDIX, consistently with previous implementations [15, 12]. Notably, *D*_*i*_ was the noisiest of all estimates, frequently hitting the upper bound of the range as shown in Supplementary Figure **??**. The challenge in estimating *D*_*i*_ with good accuracy and precision is also reported in previous NEXI and SANDI works [7, 15, 12, 14, 9].

NEXI and SMEX cortical extracellular diffusivity values are comparable between different denoising strategies and previous implementations using Connectom Scanners (*D*_*e*_ = 0.9−1 *µm*^2^/ms). However, they seem to increase when using clinical scanners (Table 1, section C). The same trend was found with SANDI, which overall reported *D*_*e*_ *≈* 0.8−0.9 *µm*^2^/ms for both real and magnitude data, also lower than previously reported in clinical data using a single, longer diffusion time (1.36 *µm*^2^/ms [11]).

All models showed consistent neurite signal fractions ranging 20-40%, also aligned with previous literature. SANDI is notably the model which estimates the lowest *f* (0.18 −0.20) in the CDMD dataset.

For Soma Fraction (*F*_*s*_), SANDI and SANDIX estimated from multiple diffusion times showed mean estimates of *≈* 0.10. These estimates were substantially lower than those previously found in [11] on a clinical scanner using a single diffusion time (*F*_*s*_ = 0.26). Here, using Δ = 19 ms only increased the soma fraction to 16%.

The soma radius (*R*_*s*_) estimate differs between SANDI and SANDIX (*R*_*s*_ *≈*24*µm* vs *R*_*s*_ *≈*17*µm*, respectively) but not between denoising strategies. When using Δ = 19 ms alone, the radius estimate decreased to *R*_*s*_ = 9.5*µm*, more in line with previous SANDI works and with biological plausible estimates.

#### 4.1.2 Quality of Fit

Figure 2 compares measured experimental data (average cortical ribbon signal) with the estimated fits from GM biophysical models. Fits are performed voxel-wise, and the resulting predicted curves are averaged across cortical ribbon voxels.

**Figure 2:**
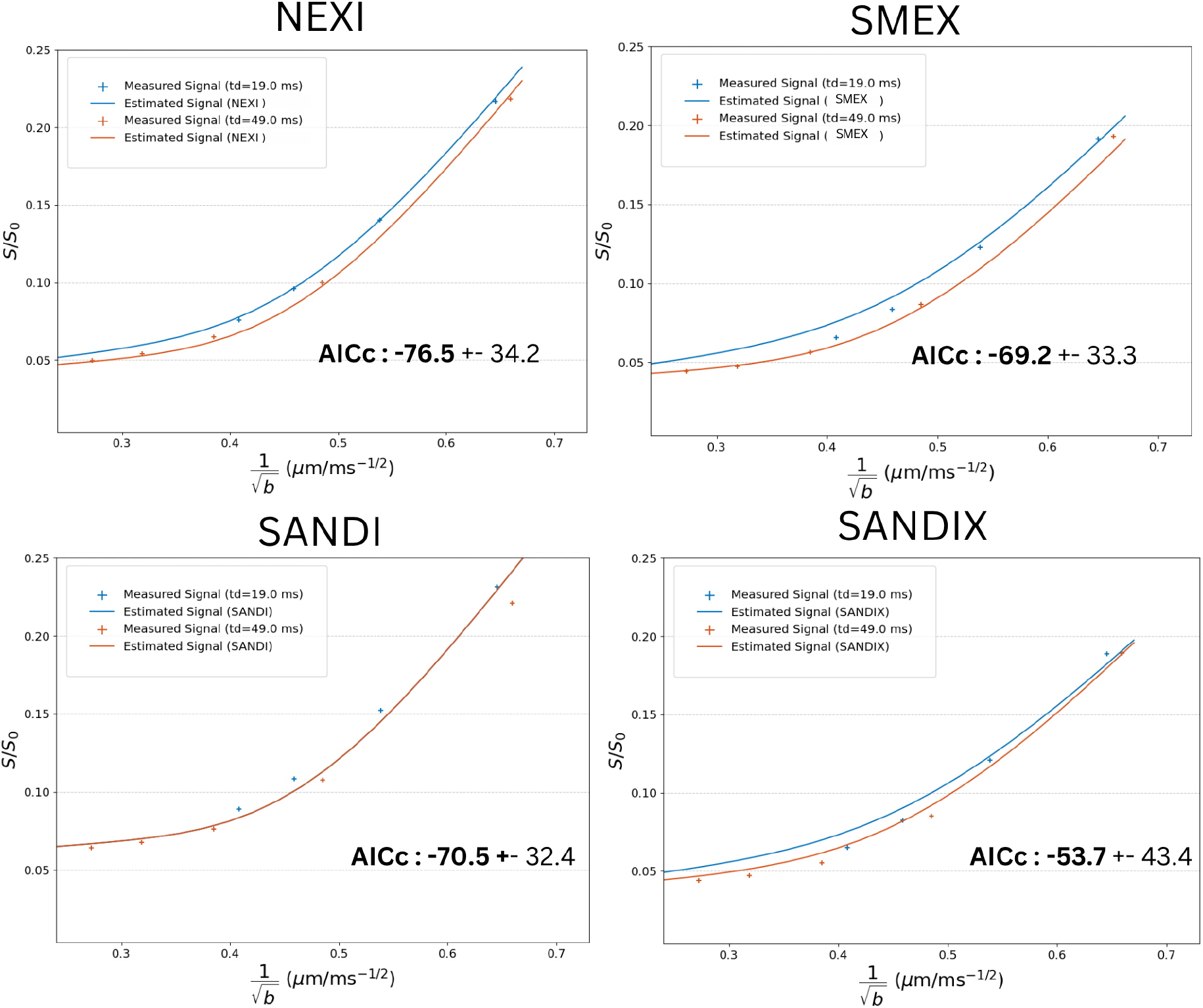
Measured vs. model-predicted signal decay across cortical ribbon voxels as a function of 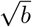 at diffusion times of 19 ms and 49 ms. Results are shown for SMEX, NEXI, SANDI, and SANDIX applied to magnitude data for all 26 subjects. Dots represent experimental signal and curves show voxel-wise model predictions averaged across the cortical ribbon. The figure highlights model accuracy and differences across diffusion times, *b*-values, and data types. Goodness of fit was assessed using AICc, with lower values indicating better fit. While 16 *b*-values were acquired in total, the plots display model fits and measured signal for the upper range 2.3 to 13.5 ms*/µ*m^2^.

**Figure 3.**
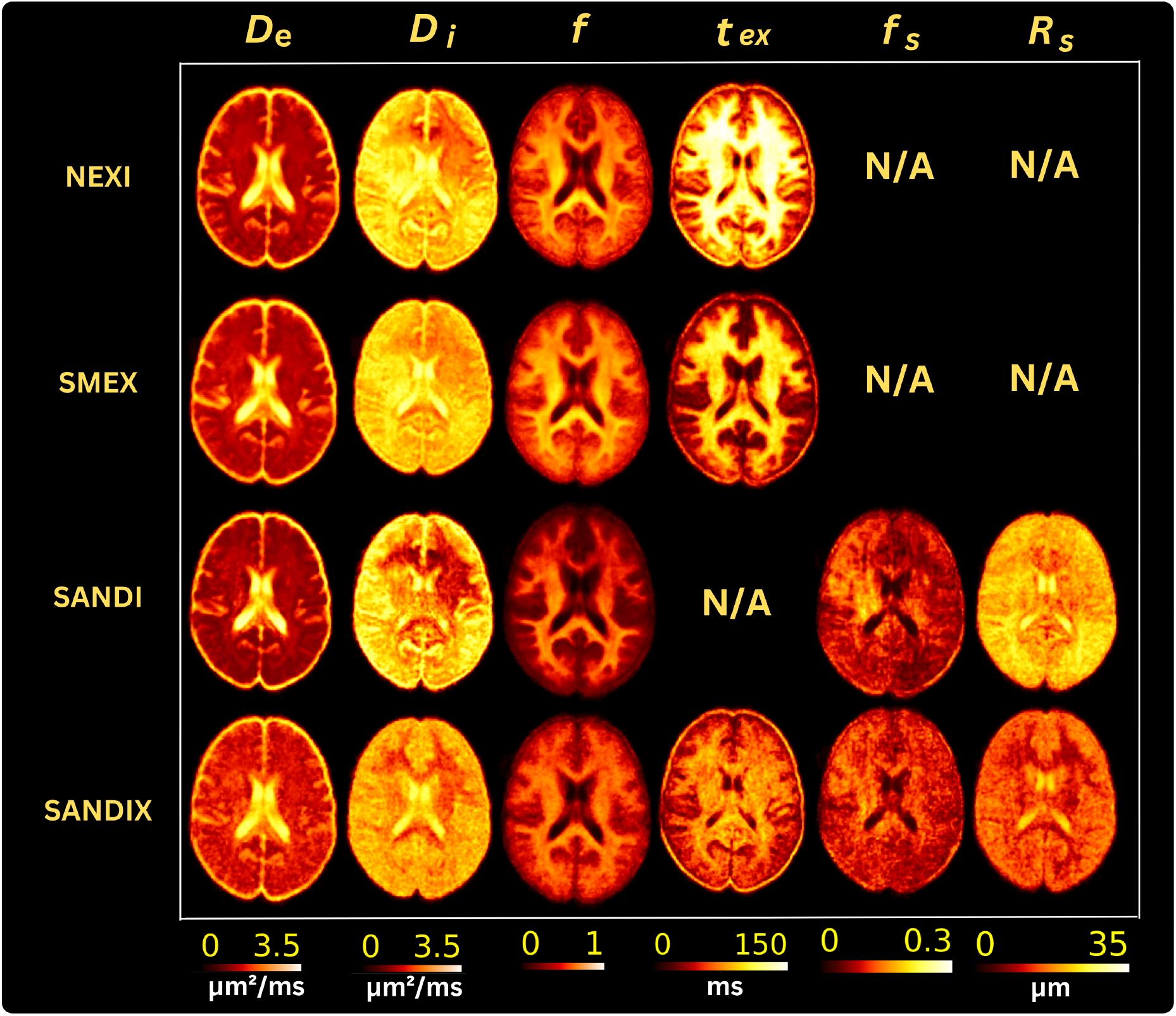
Average Parameter Maps Across Subjects in MNI Space: Axial slices of average parameter maps generated for each biophysical model, calculated across 26 subjects and represented in standard MNI space. The maps showcase spatial distributions of key model parameters, enabling a direct comparison of how different models characterize biophysical properties across the brain.

Overall, the quality of the fit illustrates that NEXI is better than SMEX, SANDI and SANDIX at fitting the measured signal (AICc= −76.5). SANDI and SMEX have similar goodness-of-fit and are followed by SANDIX. The good performance of NEXI is expected in a short gradient pulse configuration (*δ*=8 ms). Remarkably, the SMEX fit of short diffusion time data is poorer than the NEXI counterpart. SANDIX was the hardest model to fit due to the increased number of parameters to estimate, which greatly affects precision.

Nevertheless, a better fit does not always guarantee a more accurate representation of the underlying parameters, and the quality of the fit might be influenced by voxels in the cortical ribbon with partial volume effects. The examination of voxel signal distributions in the cortical ribbon (Supplementary Figure **??**) reveals that SMEX mitigates extreme values and eliminates the multi-modal distribution observed for *t*_*ex*_ and *f* with NEXI.

The comparison between real-valued and magnitude data fits is provided in Supplementary Material (Figure **??**). Consistently across all models, magnitude data processed with Rician mean correction yielded lower AICc scores than real-valued data, indicating a better fit. This improvement could be due to the incomplete removal of noise bias by the denoising method applied to real valued data.

#### 4.1.3 Group average Parametric Maps for all models

The group average parametric maps confirm the trends outlined in the ROI analysis of Table 1 and the goodness of fit in Figure 2: SANDIX maps are the noisiest across models, and *D*_*i*_ maps are the noisiest across model parameters. In SANDI, soma fraction *F*_*s*_ and radius *R*_*s*_ also display limited precision, underlying the challenge in estimating soma alongside neurite and extracellular compartments.

While it should also be noted that none of these models are designed or suited for WM microstructure, the WM-GM contrast patterns display the expected trends. Neurite density (*fn*) shows the expected higher fraction in WM vs GM. In those models that consider exchange (NEXI and SANDIX), the exchange time is also longer in WM than GM, consistent with a higher density of myelinated “impermeable” axons and thus a slower inter-compartment exchange. Finally, the estimated soma fraction *F*_*s*_ and radius *R*_*s*_ are higher in the cortex than in WM, which aligns with the biological representation that cellular bodies (soma) occupy approximately 10–20% of GM by volume and are not as present in WM.

For all models and all parameters except for *D*_*i*_ and *F*_*s*_, the axial maps show an expected degree of left-right symmetry between hemispheres. The unexpected asymmetry in *D*_*i*_ and *F*_*s*_ could be due to residual field gradient inhomogeneities, that are captured by the model parameters with highest uncertainty.

### 4.2 Focused Comparison on NEXI vs SMEX

#### 4.2.1 Simulation

Estimation errors on synthetic SMEX data revealed differences in fitting performance between the SMEX and NEXI models (Figure 4). *D*_*e*_ and *f* showed good accuracy and precision for both model variants, while *t*_ex_ and *D*_*i*_ were less precise, consistent with previous reports [15, 12].

**Figure 4.**
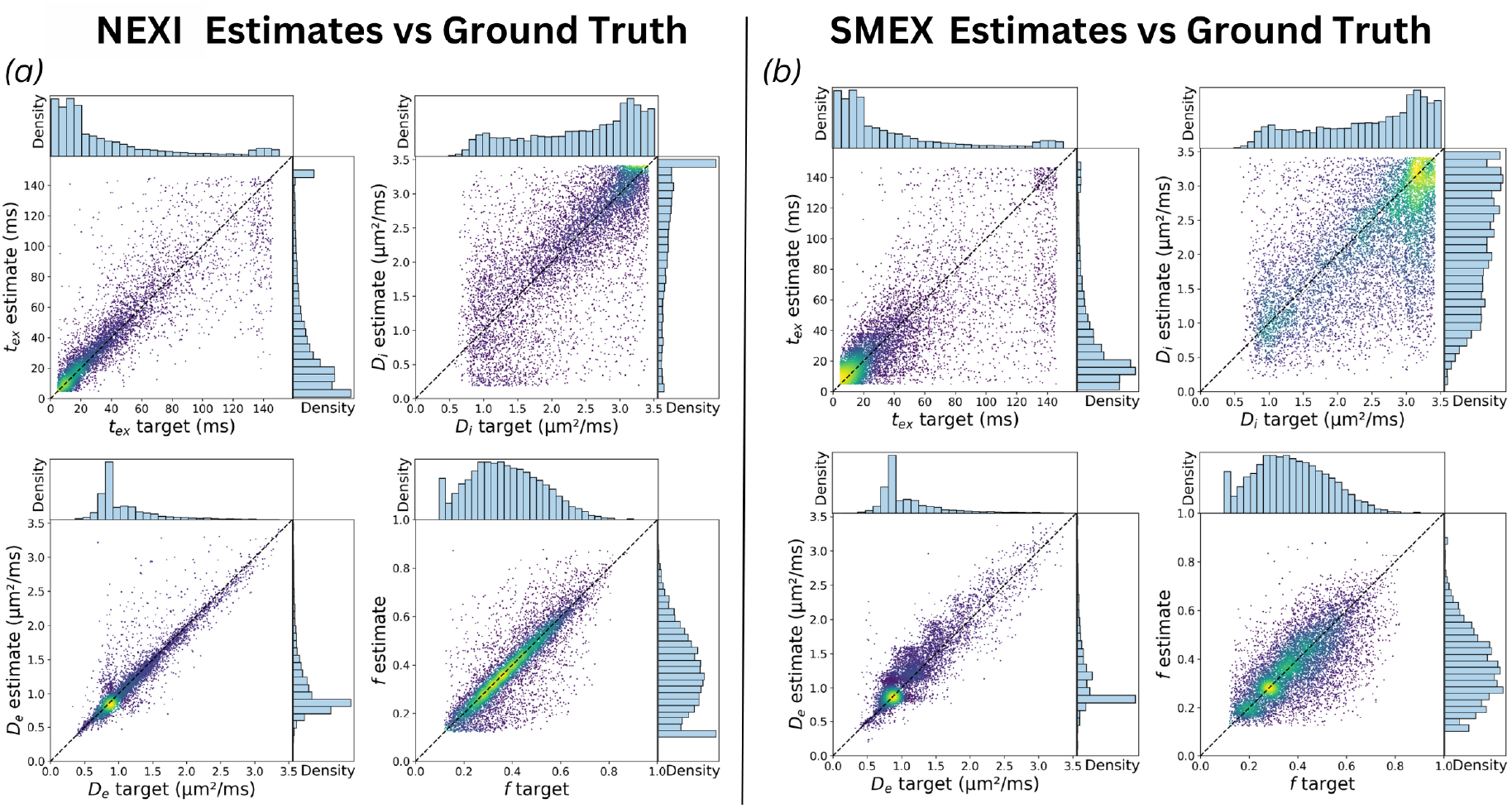
Comparison of parameter estimation from synthetic SMEX signals. Density-colored scatter plots show estimated vs. ground truth parameters (*t*_ex_, *D*_*i*_, *D*_*e*_, *f*) for NEXI (a) and for SMEX (b) Synthetic ground truth SMEX signals were generated using the same *δ*, diffusion times, and *b*-values as in vivo acquisitions. Signals were then powder-averaged to emulate magnitude data. Histograms show marginal distributions; the dashed diagonal indicates perfect estimation.

Despite being fitted to the same noisy synthetic signals, surprisingly, NEXI showed better estimation performance across parameters. In contrast, SMEX exhibited larger deviations in several parameters, particularly for *f*, *D*_*e*_, and *t*_ex_.

SMEX preserves better the ground truth distribution of *D*_*i*_ values, while NEXI estimates tend to aggregate towards the upper bound, reflecting poor specificity.

#### 4.2.2 Surface Maps for NEXI and SMEX

Figure 5 shows human cortical surface maps for NEXI and SMEX estimated using two diffusion times. As expected, the two models show almost identical anatomical trends, with slight variations in their values and thus scales. SMEX shows lower *t*_*ex*_, *f*, *D*_*e*_, and *D*_*i*_ than NEXI, which is consistent with other works [12].

**Figure 5.**
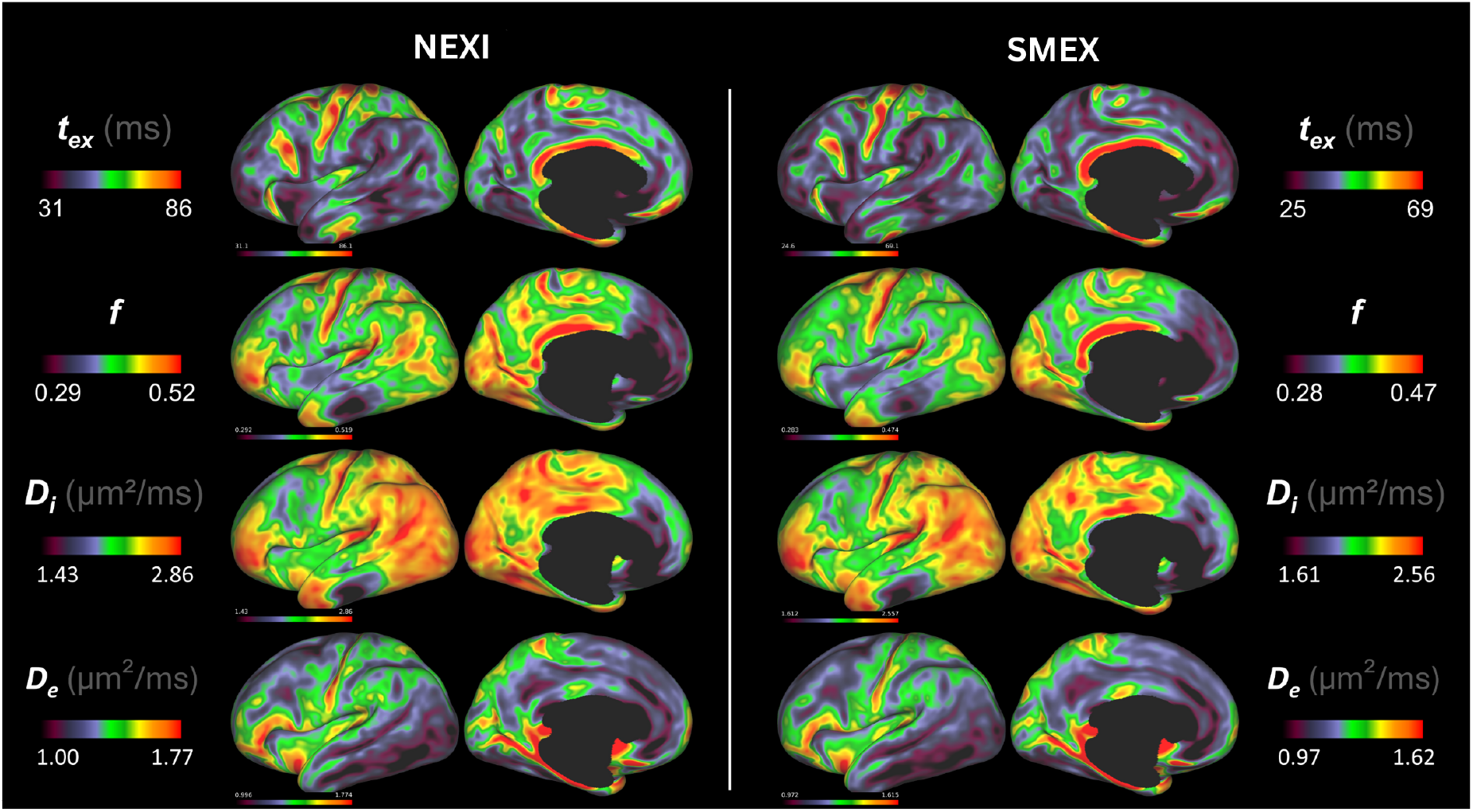
Projection onto cortical surface in fsaverage space of NEXI and SMEX, maps averaged across 25 subjects. Maps show a slight shift in the scales between models, but both show very similar anatomical trends.

The estimated intercompartmental exchange time (*t*_ex_) exhibits elevated values in the somatosensory and motor cortices, including the precentral and paracentral regions, as well as in the occipital lobe and temporal lobe.

Similarly, the neurite volume fraction (*f*) demonstrates higher values in the occipital lobe. The relatively lower *f* observed in the anterior temporal lobe is also consistent with findings from NEXI studies conducted on clinical MRI systems [12].

Although the intra-axonal diffusivity (*D*_*i*_) map is the noisiest of the estimated parameters, it reveals some anatomical patterns that align with biological and model expectations. Specifically, *D*_*i*_ appears to follow a similar spatial distribution to *f*, reflecting the association that regions with higher neurite density would result in faster intra-neurite diffusivity.

Finally, the estimated extra-axonal diffusivity (*D*_*e*_) exhibits spatial distributions consistent with prior NEXI studies on both Connectom [15] and clinical scanners [12]. Notably, *D*_*e*_ shows reduced values in the occipital lobe and elevated values in the somatosensory cortex. In addition, we observed faster *D*_*e*_ in the insula, which was also reported previously [15].

## 5 Discussion

This analysis compared parameter estimates from four biophysical models using an open-source CDMD dataset [13] to quantify different microstructural properties of cortical tissue in the human brain. Here, the newly developed processing package “Gray Matter Swiss Army Knife” [10] processed NEXI [7], SMEX [8], SANDIX [8] and SANDI [9] using two diffusion times instead of the four, typically employed in earlier implementations of exchange models. This study serves as a proof of concept to assess whether these models can be reliably estimated from only two diffusion times. It compares parameter estimates across models, subjects, and previous studies, and assesses the quality of the model fits to experimental data.

Previous NEXI papers have highlighted the importance of accounting for the Rician mean floor either in signal model or data processing for accurate fitting of NEXI parameters (particularly *t*_ex_) [15, 12]. Therefore, we investigated how two different denoising strategies impact parameter estimation across all four biophysical models: (1) Complex Denoising reduces Gaussian noise from the raw complex-valued data to then retain the real-valued signals in the fitting procedure; (2) Magnitude Denoising reduces noise in magnitude data and uses the concomitant noise variance estimation to correct for Rician bias in the model fit.

In addition, since NEXI and SMEX share the same underlying microstructural assumptions but differ in their mathematical formulations, we performed a focused comparison between them. Specifically, we evaluated (1) their accuracy in recovering synthetic ground truth signals using simulations, and (2) the anatomical plausibility of their cortical surface maps in the absence of a true reference.

CDMD data fits showed that parameters common to all four models,namely *f*, *D*_*i*_, and *D*_*e*_, yielded cortical ribbon metrics in the ranges of *f* ~ 0.19–0.32, *D*_*i*_ ~2.2–3.0 *µ*m^2^*/*ms, and *D*_*e*_ ~0.79–1.09 *µ*m^2^*/*ms. The estimated neurite fraction *f* was lower in SANDI and SANDIX compared to NEXI and SMEX. This likely reflects the explicit modeling of a soma compartment, which redistributes part of the intracellular signal from neurites to soma. As a result, the total intracellular fraction (*f* + *F*_*s*_) remains *~* within a similar range (~ 20–40%). This finding, however, contrasts with the common interpretation in NEXI and SMEX that the soma contribution is pooled into the extracellular compartment. Neurite fraction estimates for all models are in line with previous *in vivo* dMRI studies in both rodents and humans, which consistently report cortical GM neurite fractions in the range of 20–40% [7, 15, 12,14, 9, 30, 31, 8]. By contrast, histological studies often report higher neurite fractions, up to 60% [32], though this may reflect fixation-induced shrinkage of the extracellular space in ex vivo tissue. Supporting this, ex vivo NEXI studies report higher neurite fraction estimates, closer to the histological findings [8, 33].

Consistently across this and previous studies, *D*_*i*_ was found to be the least robust parameter for all models, often exceeding 2.5 *µ*m^2^*/*ms and frequently reaching the upper fitting bound. Estimated *D*_*i*_ values across models and for both magnitude and real-valued data ranged between 2.20 and 2.94 *µ*m^2^*/*ms, at times barely lower than the free water diffusivity at body temperature (3 *µ*m^2^*/*ms) [34]. Finally, parameter maps reveal substantial noise, underscoring the instability of this parameter, likely due to a flat fitting landscape along this dimension of parameter space [35]. Among the tested models, SANDIX and SMEX yielded the most biologically plausible *D*_*i*_ estimates, aligning better with axonal diffusivity values reported in WM [26, 36], with *D*_*i*_ values ranging 2.20 to 2.46 *µ*m^2^*/*ms. It is noteworthy that both these models account for the actual pulse duration *δ*.

Previous works have shown that multi-diffusion time datasets carry the signature of exchange rather than of soma restriction, in the form of lower signal with longer diffusion times, at a fixed b-value [7, 8, 12]. Two diffusion time data are expected to affect SANDI estimates substantially because the soma restriction would predict increasing signal with longer diffusion times, while the experimental data display the opposite behavior. As a result, the predicted signals from SANDI overlap for the two diffusion times in the case of magnitude data, or their order is swapped (longer diffusion time predicted to have higher signal than shorter diffusion time) in real-valued data (Supplementary Figure **??**). A recent study [24] using real-valued data from the same CDMD dataset compared SANDI estimates at Δ = 19 ms, Δ = 49 ms, and the SANDIX model, revealing a strong underestimation of neurite signal fraction at longer diffusion times. This effect was attributed also to inter-compartmental water exchange not being accounted for, while becoming more prominent as diffusion time increases.

Our data shows SANDI estimates from the joint fit of both diffusion times differed quite notably from previous SANDI reports of the human cortex, whether from a Connectom 1 scanner [29, 37] or a clinical scanner [11]. As expected, when fitted with one diffusion time (Δ = 19 ms) SANDI estimates were more aligned with previous reports. The soma fraction increased from 10% to 16%, and the estimated soma radius *R*_*s*_ decreased from 24.3 *µ*m to 9.5 *µ*m, which are much more aligned with other imaging literature [30, 38, 16]. A recent morphological study that analyzed 3,500 GM cells estimated that neural soma sizes range from 2 to 30 *µ*m in radius with an average of 7 *µ*m [39]. Although still falling within the suggested range, there is a probable overestimation of the radius in these biophysical models, due to the volume weighting and the indirect non-linear probe via diffusivity. A previous study estimated that this overestimation could be up to a factor of 1.6 for soma radii [8].

The goodness of fit analysis showed that NEXI was the best at fitting parameter estimates to powder-averaged signal in the cortex. Similarly, SANDI fits suggest the exchange term is missing from the model, due to the challenge of capturing the correct time-dependence. To address these limitations, more advanced GM models are currently being explored, such as SANDIX or the Generalized Exchange Model (GEM), which introduces a dedicated soma compartment along with exchange dynamics within the soma, aiming for improved biological accuracy [40].

Consideration of other components, like myelinated axons in GM, have also been previously proposed – enhanced SANDIX (eSANDIX) [8]. Although the signal contribution of myelinated axons is relatively small, it may become non-negligible at high *b*-values and could introduce subtle biases in parameter estimation if not modeled appropriately.

However, this increased biophysical realism comes at the cost of model complexity and a higher number of free parameters, which can hinder parameter stability. Indeed, SANDIX shows the worst fit and noisiest parameter maps. Therefore, careful consideration is needed when extending biophysical models, especially when applied to in vivo data with inherent limitations in signal-to-noise and acquisition duration. Currently, the trade-off between biological realism and model stability remains a central challenge in GM diffusion modeling. Further research into the contributions of diffusion signal from different components should be conducted to improve these models and balance specificity with precision.

Both NEXI implementations, designed to capture exchange between neurites and the extracellular space and typically requiring multiple diffusion times, produced parameter estimates comparable with previous studies that used four diffusion times.

In this dataset too, correction for the Rician floor in the magnitude data yielded lower *t*_ex_ than NEXI without Rician mean correction (albeit on real-valued data). The Rician mean correction has been shown to reduce estimation bias and stabilize *t*_ex_ given its high susceptibility to the noise floor [15]. A recent study [14] using NEXI reported that estimation bias was reduced by a factor of ≈ 1.5 when using NEXI-RM compared to ordinary NEXI. Moreover, it was shown that this correction had impact on the Connectom 1.0 scanner, but not on the Connectom 2.0, where the SNR is high enough for the Rician floor to be negligible.

Moreover, despite the use of phase images for complex denoising, residual noise still appears to influence the real-valued data and the resulting *t*_ex_ estimation, reaffirming the need for explicit noise modeling even in real-valued fitting. We observe that *t*_ex_ values derived from real-valued data are substantially higher, show larger confidence intervals (particularly for SMEX), and exhibit poorer goodness-of-fit. It is therefore recommended to implement a Rician mean correction on magnitude data when fitting either NEXI or SMEX, unless ultra-strong gradients that can yield high SNR images, such as those in Connectom 2.0, are available.

The better performance of NEXI in terms of goodness-of-fit and accuracy and precision in simulations is somewhat unexpected, as prior work had shown SMEX to outperform it on synthetic signals. However, this difference is likely attributable to numerical implementation factors rather than fundamental model properties. Specifically, the solver used in SMEX relies on the numerical integration of a differential system via scipy.integrate.solve ivp, which introduces additional imprecision — particularly problematic when high numerical precision is needed. In contrast, NEXI involves a direct integration using scipy.integrate.quad vec, which tends to yield more stable results. Furthermore, in previous studies, the advantage of SMEX was shown when the NPA was indeed not met (e.g. with *δ* = 16.5 ms), making NEXI less appropriate. In this work, with a shorter pulse duration (*δ* = 8 ms), the assumptions behind NEXI are better met, and the numerical stability of its solver likely further contributes to its improved performance.

The anatomical patterns observed in our surface maps, particularly for parameters such as exchange time (*t*_ex_) and neurite fraction (*f*), are consistent with previously reported NEXI implementations and with known cytoarchitectonic and myeloarchitectonic cortical features [41]. Specifically, the elevated *t*_ex_, *f* and *D*_*i*_ values in primary sensorimotor cortices (including precentral, postcentral, and paracentral regions) align with the dense myelination and cell packing reported in classical histological and post mortem high field MRI studies [42]. These regions are known to contain large, heavily myelinated pyramidal cells (e.g., Betz cells in M1) [43], which likely contribute to the prolonged exchange times due to reduced permeability of their membranes.

The higher *f* observed in the occipital lobe, particularly in the primary visual cortex (V1, Brodmann area 17), is consistent with histological findings showing densely packed vertical bundles of apical dendrites [44, 41], as well as with previous imaging studies [15, 12, 9, 31]. Conversely, the relatively lower *f* observed in the anterior temporal lobe aligns with cytoarchitectonic descriptions of reduced cell density and laminar differentiation in this region [44], as well as with more recent parcellation studies based on microstructural imaging [45].

Furthermore, the surface map distribution of intra-axonal diffusivity (*D*_*i*_), although noisier, appears to follow similar patterns as *f* with a different scale, suggesting that regions with denser neurite packing naturally present faster intra-neurite diffusivity. Indeed, higher neurite density may go in pair with more straight dendrite segments to reach a high packing, which in turn yields faster *D*_*i*_. Meanwhile, extra-axonal diffusivity (*D*_*e*_) shows a distinct pattern with lower values in the occipital and temporal lobes and higher diffusivity in the somatosensory and insular cortices. These patterns have also been observed in prior NEXI studies using Connectom 1.0 protocols [15].

Together, these results highlight that the CDMD dataset-derived parameter and surface maps reflect known biological distributions of cortical microstructure and are consistent across other NEXI and SMEX implementations, other biophysical models and neuro-anatomical patterns from other imaging modalities. Importantly, they support the feasibility of probing gray matter microstructure using only two diffusion times, offering a time-efficient alternative to conventional protocols that typically require three or four diffusion times for NEXI estimation.

It is important though to highlight that, although the diffusion MRI protocol in this study used only two diffusion times, it included a total of 16 *b*-values, 32 to 64 diffusion gradient directions depending on the *b*-value, with a total scan duration of 55 minutes. While this is shorter than some high-angular resolution protocols, this overall acquisition remains lengthy and limits feasibility in clinical environments due to its complexity.

Future work will focus on reducing scan time while preserving parameter sensitivity. Optimizing the combination of diffusion times, *b*-values, and gradient directions could enable development of a more time-efficient protocol, improving applicability in both research and clinical settings [46, 47].

## 6 Conclusions

We retrospectively evaluated and compared the performance of four advanced biophysical models, NEXI, SMEX, SANDI and SANDIX, on a publicly available diffusion MRI dataset (CDMD) [13] fitted with the Gray Matter Swiss Knife Package [10].

These models varied in their goodness of fit against experimental data, with NEXI showing the best fit, followed by SANDI and SMEX, and finally SANDIX.

The parameter estimates obtained in this study are consistent with those reported in previous studies. For exchange models such as NEXI and SMEX, we demonstrate that accurate and anatomically meaningful parameter estimates can be obtained using an acquisition protocol with only two diffusion times. Crucially, we emphasize the importance of explicitly modeling the noise, even when processing real-valued data, as it significantly affects the exchange time parameter estimates (*t*_ex_). In the case of magnitude data, this translates into incorporating the Rician mean in the NEXI model, as previously introduced [15]. Additionally, we show that the SANDI model performs best when fitted using a single short diffusion time. Including multiple diffusion times degrades the soma radius estimation due to inadequate modeling of time dependence, while using only a long diffusion time leads to underestimation of the neurite fraction.

Future efforts should focus on further optimizing acquisition parameters to reduce the scan time and enable the clinical translation of GM models to patient populations, while ensuring interpretability and reliability of cortical microstructure estimates.

## 7 Acknowledgments

This work was supported by the Swiss National Science Foundation under an Eccellenza Fellowship no. 194260 (to I.J.), by the University of Lausanne and the Lausanne University Hospital (CHUV).

