## Supplementary Materials for "Comparative Systematic Analysis of Gray Matter Biophysical Models on a public dataset"

### Supplementary Material

#### MGH Connectome Diffusion Microstructure Dataset (CDMD) Acquisition Protocol

**Imaging Parameters:**

- *Repetition Time (TR)*: **3800 ms.**
- *Echo Time (TE):* **77 ms.**
- *Field of View (FOV)*: **216 × 216 mm.**
- *Matrix Size*: **108 × 108.**
- *Slice Thickness:* **2 mm.**
- *Voxel Size:* **2 × 2 × 2 mm^3.**
- *Diffusion Times (Δ):* **19 ms or 49 ms.**
- *Diffusion-Encoding Gradient Duration (δ)*: **8 ms.**

**For Δ = 19 ms:**

- **B-values:**
  - 50 s/mm$^2$ - 32 directions
  - 350 s/mm$^2$ - 32 directions
  - 800 s/mm$^2$ - 32 directions
  - 1500 s/mm$^2$ - 32 directions
  - 2400 s/mm$^2$ - 64 directions
  - 3450 s/mm$^2$ - 64 directions
  - 4750 s/mm$^2$- 64 directions
  - 6000 s/mm$^2$ - 64 directions

**For Δ = 49 ms:**

- **B-values:**
  - 200 s/mm$^2$ -32 directions
  - 950 s/mm$^2$ - 32- directions
  - 2300 s/mm$^2$ - 32 directions
  - 4250 s/mm$^2$ - 64 directions
  - 6750 s/mm$^2$ - 64 directions
  - 9850 s/mm$^2$ - 64 directions
  - 13,500 s/mm$^2$ - 64 directions
  - 17,800 s/mm$^2$ - 64 directions
- **Diffusion Encoding Directions:**
  - 32 uniform directions for b-values < 2400 s/mm$^2$.
  - 64 uniform directions for b-values ≥ 2400 s/mm$^2$.

Figure S1: CDMD acquisition parameters.

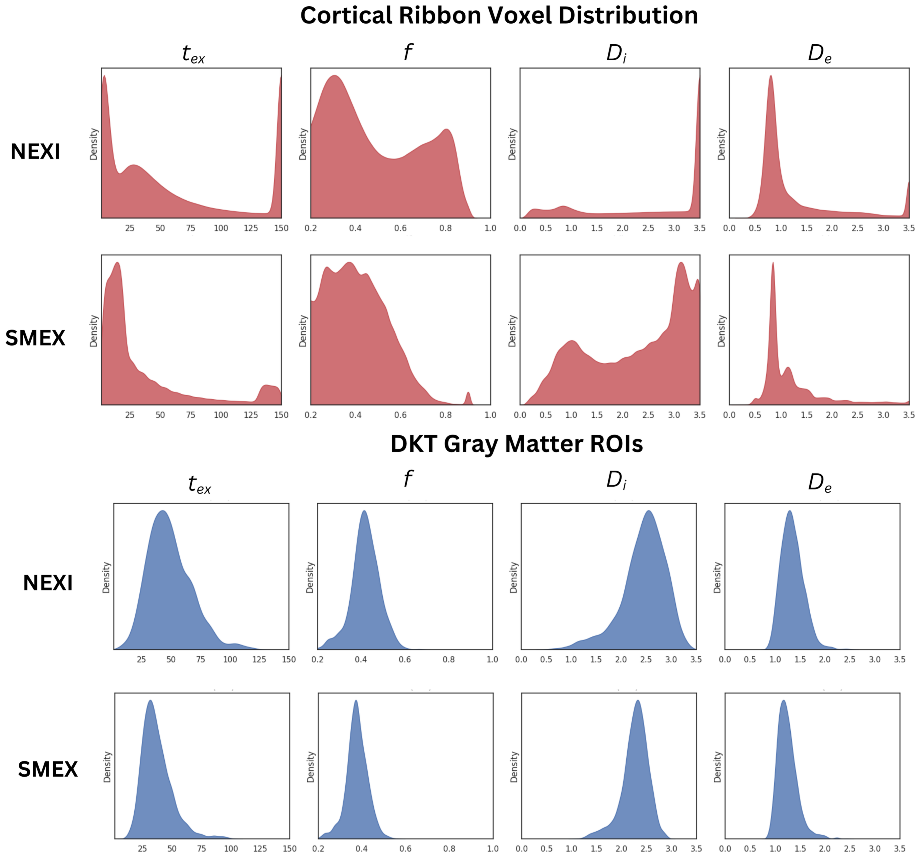


Figure S2: Distributions of voxel-wise parameter estimates and DKT ROI-wise medians in the cortical ribbon for NEXI and SMEX models, using magnitude data from the 26 subjects. These distributions closely resemble those reported in [?] (Figure 3), notably showing that NEXI tends to produce values near the fitting bounds and exhibits a bimodal distribution of the neurite fraction $f$, which is not present in SMEX. Conversely, SMEX displays comb-like peaks in certain parameters, likely due to interactions between the ODE-based signal model and the initial grid search.

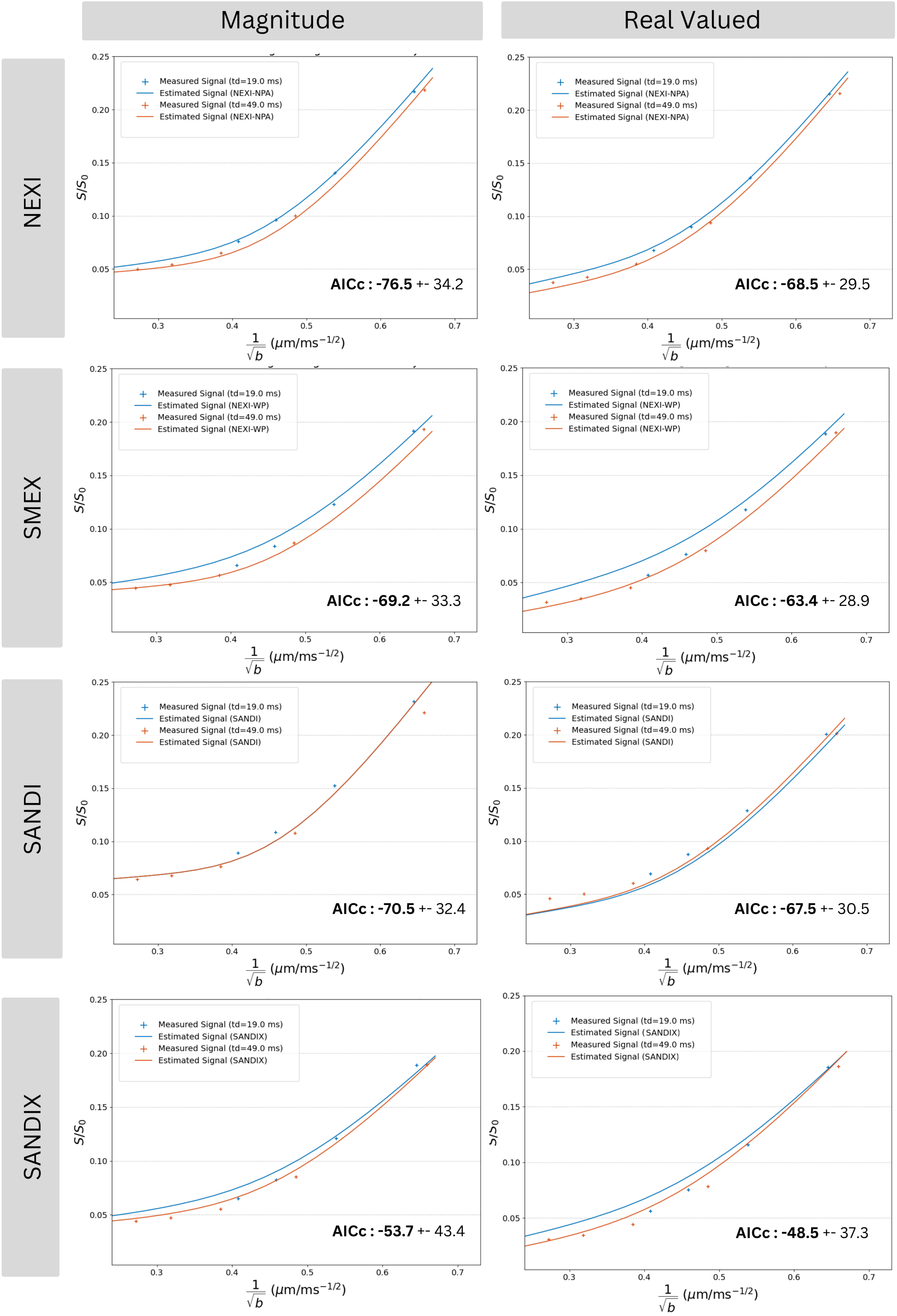


Figure S3: Quality of fit comparison between magnitude data (26 subjects) and real-valued data (14 subjects). Dots represent experimental signal and curves show voxel-wise model predictions averaged across the cortical ribbon. The figure highlights model accuracy and differences across diffusion times, $b$-values, and data types. Goodness of fit was assessed using AICc, with lower values indicating better fit.
